# Mathematical Model of Muscle Wasting in Cancer Cachexia

**DOI:** 10.1101/2020.03.30.016709

**Authors:** Suzan Farhang-Sardroodi, Kathleen P. Wilkie

## Abstract

Cancer cachexia is a debilitating condition characterized by an extreme loss of skeletal muscle mass which negatively impacts patient’s quality of life, reduces their ability to sustain anticancer therapies, and increases the risk of mortality. Recent discoveries have identified the myostatin/activin-ActRIIB pathway as critical to muscle wasting by inducing satellite cell quiescence and increasing muscle-specific ubiquitin ligases responsible for atrophy. Remarkably, pharmacological blockade of the ActRIIB pathway has shown to reverse muscle wasting and prolong the survival time of tumor-bearing animals. To explore the implications of this signaling pathway and potential therapeutic targets in cachexia, we construct a novel mathematical model of muscle tissue subjected to tumor-derived cachexic factors. The model formulation tracks the intercellular interactions between cancer, satellite cell, and muscle cell populations. The model is parameterized by fitting to colon-26 mouse model data, and analysis provides insight into tissue growth in healthy, cancerous, and post-treatment conditions. Model predictions suggest that cachexia fundamentally alters muscle tissue health, as measured by the stem cell ratio, and this is only partially recovered by anti-cachexia treatment. Our mathematical findings suggest that the activation and proliferation of satellite cells, after blocking the myostatin/activin B pathway, is required to partially recover cancer-induced muscle loss.

## 1. Introduction

Cancer cachexia is a condition identified by an ongoing and irreversible loss of skeletal muscle and adipose tissue (1–4). Depending on the tumor type, cachexia affects 30 to 80% of cancer patients (5) and it leads to increased morbidity and mortality (6–8). The highest incidence of cachexia is seen in patients with pancreatic, gastric or lung cancers (9, 10). Historically, the cause of cancer cachexia was thought to be metabolic dysregulation (9, 11), which led to nutritional and fitness-based therapeutic approaches. None of these approaches were successful, however, leaving cachexia as an untreatable condition that severely impacts patient quality of life (12–14). Extreme weight loss is the primary clinical feature by which cachexia is defined. Generally, a body weight drop of more than 25% is considered cachexia (1). The dramatic weight loss causes increased disease symptoms such as pain, weakness, and fatigue (15), and leads to limited treatment options (16, 17).

Recently, a new paradigm arose in cancer cachexia research, proposing that tumor-derived factors may alter tissue behavior through molecular signaling. Various tumor-derived factors such as cytokines, hormones, and glucocorticoids can influence muscle protein balance through signaling transduction pathways (18). From these various factors, the myostatin/activin A signaling was found to play a significant role in cancer-induced muscle atrophy (19–23). Indeed, blocking the activin type-2 receptor (ActRIIB) with a soluble decoy receptor (sActRIIB) or a myostatin antibody in several mouse models of cachexia has been shown to prevent muscle wasting, and also to reverse the prior loss of skeletal muscle (21, 24–27). Consequently, the survival time of the tumor-bearing animals was increased, even though the tumor itself was not therapeutically targeted.

Myostatin and Activin *A* are members of the transforming growth factor-*β* (TGF-*β*) superfamily of proteins. They both bind to ActRIIB, a high affinity receptor (28–31), resulting in the phosphorylation of Smad2 and dephosphorylation of FOXO3a which consequently induces satellite cell quiescence. The dephosphorylated FOXO3a then moves into the nucleus and induces two muscle-specific ubiquitination ligases that facilitate the degradation of myofibrillar proteins, leading to muscle loss and cachexia (10, 21, 32–35).

Recent work by Zhou *et al*.(21) demonstrates that blocking the activation of ActRIIB and the subsequent downstream signaling events can stop muscle loss in cancerous mice. They assessed the effects of an ActRIIB decoy receptor in several mouse models of cancer cachexia, including a murine colon-26 carcinoma and two xenograft models: human G361 melanoma and human TOV-21G ovarian carcinoma. The decoy receptor sActRIIB potentially prevents binding of myostatin and activin A to the ActRIIB receptor and inhibits the downstream signaling. In each case, treatment with sActRIIB was found to prevent or reverse skeletal muscle atrophy and heart muscle wasting, although it had no effect on adipose tissue loss or the production of pro-inflammatory cytokines (for colon-26-bearing mice).

Under normal conditions, adult skeletal muscle preserves its mass via feedback mechanisms that balance the signals controlling muscle catabolism and anabolism. When signaling is dysregulated by tumor-derived factors, the implications on tissue homeostasis are not clear due to the nonlinear and complex dynamics involved in such feedback control mechanisms. Therefore a quantitative tool is needed to better understand the significant factors regulating muscle atrophy (increased muscle protein degradation) and hypertrophy (increased muscle protein synthesis). Mathematical modeling is one such tool, and the predictive nature of the model presented in this paper can be used to examine the complex nature and potential treatment of cancer cachexia.

We present a novel mathematical framework to explore the effects of dysregulated signaling in cancer cachexia. Inspired by a model for population growth with negative feedback (36), we construct a model of feedback-regulated healthy muscle tissue by describing the growth rates of muscle and satellite cells (muscle stem cell) compartments using ordinary differential equations. Negative feedback from the muscle to stem compartments maintains tissue homeostasis. We then extend the model to include a growing tumor that interferes with the intercellular signals, and thus, with the feedback control. This dysregulation leads to muscle loss in the model. Finally, the model is extended to simulate the effects of ActRIIB blockade treatment. Model parameters are found by fitting to experimental data for healthy, diseased and recovery states, and the model is validated by comparing to a second experimental group(21). Model simulations suggest that satellite cell reactivation is the primary target for muscle regulation through the myostatin/activin-ActRIIB signaling.

## 2. A Model of Healthy Muscle Tissue

Skeletal muscle is a form of striated muscle tissue, accounting for about 40% of a healthy person’s body weight (37). Adult skeletal muscle, which is stable under normal circumstances, has an extraordinary ability to regenerate post-injury by activating various precursor cells. Here we focus on the dynamics of satellite cells (the principal skeletal muscle stem cells (38)) and myofibers (the terminal differentiated muscle cells). Satellite cells comprise approximately 3% − 7% (39–42), of all muscle nuclei in skeletal muscle and are mitotically dormant unless activated by an external signal. Upon receiving an activation signal, a satellite cell will enter the cell cycle to proliferate and differentiate into myogenic precursor cells in an attempt to repair the damage and maintain tissue homeostasis (43–48).

To model the dynamics of skeletal muscle regeneration and repair, we begin with a classic model of cell lineage (36, 49). Neglecting the intermediate precursor cells to avoid unnecessary complexity, we consider only two cell compartments: the satellite cells (or stem cells) *S*(*t*), and the muscle cells (or differentiated cells) *M* (*t*):

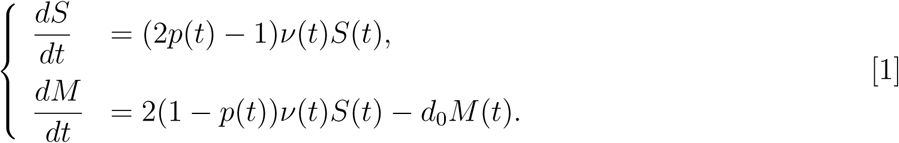

Here, *d*_0_ is the natural death rate for differentiated muscle cells. The proliferation rate of satellite cells is *ν*(*t*), and the probability of symmetric self-renewing division is *p*(*t*). That is, once activated, a satellite cell will divide into two daughter satellite cells with probability *p*(*t*), and into two differentiated muscle cells with probability 1 − *p*(*t*). The probability of self-renewal, *p*(*t*), and the proliferation rate, *ν*(*t*), are determined by intercellular interactions and thus provide the mechanism for negative feedback control. The muscle compartment will decrease the proliferation rate and probability of self-renewing division as it approaches its homeostatic level. Following muscle injury, satellite cells undergo extensive proliferation to self renew and differentiate into muscle cells (50). After resolution of the healing process, the division rate and probability return to their homeostatic maintenance levels. To model feedback in the symmetric division probability, we propose:

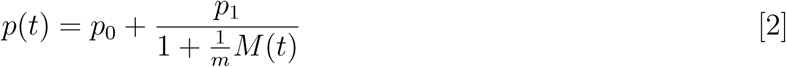

where *p*_0_ is the homeostatic probability, *p*_1_ is the perturbation probability activated in growth or in response to injury, and *m* is the half-saturation constant. Since this is a probability, we must have 0 *< p*_0_ + *p*_1_ *≤* 1. Notice from Eq. (1) that if *ν*(*t*) *>* 0 then 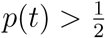 will result in unbounded growth in the stem compartment, whereas 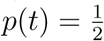 will result in no growth. We thus expect 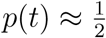 in the steady state.

Similar to the above, we propose the same feedback form for the proliferation rate of satellite cells:

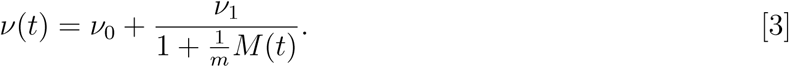

Here *ν*_0_ is the homeostatic division rate, *ν*_1_ is the perturbation division rate, activated in growth or in response to injury, and *m* is the half-saturation constant.

These negative feedback loops provide two mechanisms by which the satellite cell behavior is controlled by the muscle compartment. Together, they act to regulate the tissue to maintain homeostasis.

### Model Parameterization

The experimental data that is available to us comes in many forms: body or tissue mass in grams (g), tumor volume in mm^3^, and cell numbers (from tumor implantation). We thus need to manipulate the data to match our mathematical model description, which is nominally a volume measurement for each compartment. To parameterize our model we deconstruct body weight measurements into an estimate of lean muscle mass. We assume that body weight (mass) consists of 40% skeletal lean mass and 60% other (including adipose tissue, bone, organ tissue, etc). From this 40%, we estimate that 95% is due to muscle cells, and 5% is related to satellite cells in adult muscle. Finally, to convert mass in grams to tissue volume in mm^3^, we estimate a conversion factor of *ξ* = 0.002 g/mm^3^ (from an estimated tumor volume of 1000 mm^3^ weighing 2 g (21)[Fig 2C]). These breakdowns are estimates that allow us to fit our model with experimental data.

**Fig. 1.**
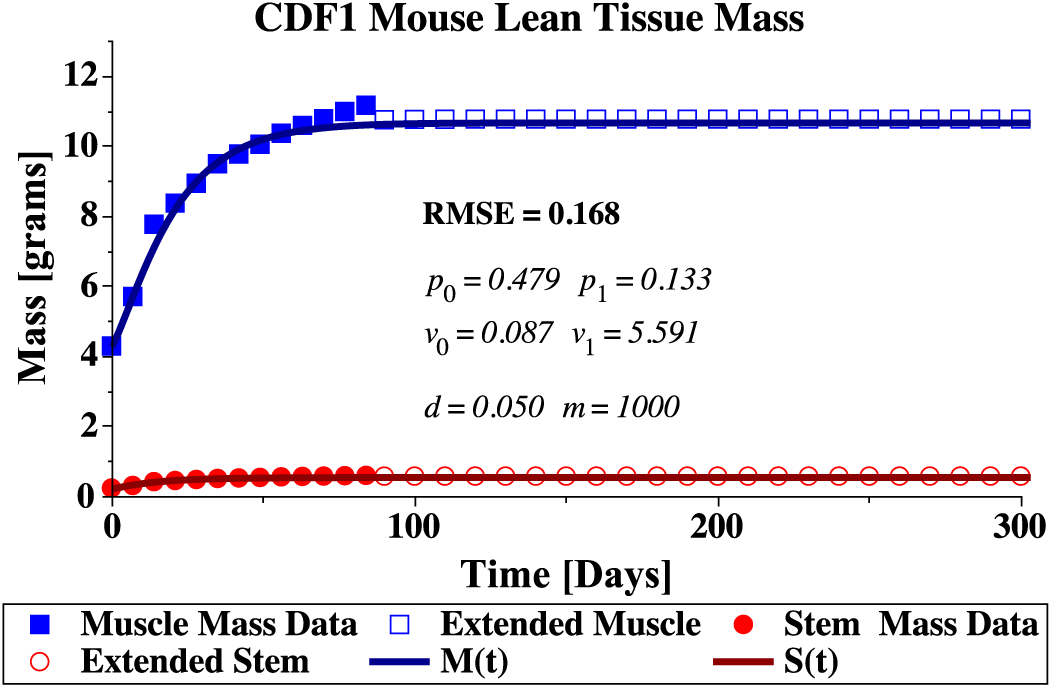
Lean body mass for a US-bread CDF1 male mouse. Body weight measurements from the suppliers published growth data (51) are deconstructed into estimated muscle (solid box) and satellite (solid circle) masses, and then extended by a logistic growth model (extended muscle mass open box and extended satellite mass open circle). Finally the estimated and extended data points are used to fit the mathematical model given by equations Eq. (1)–Eq. (3).

**Fig. 2.**
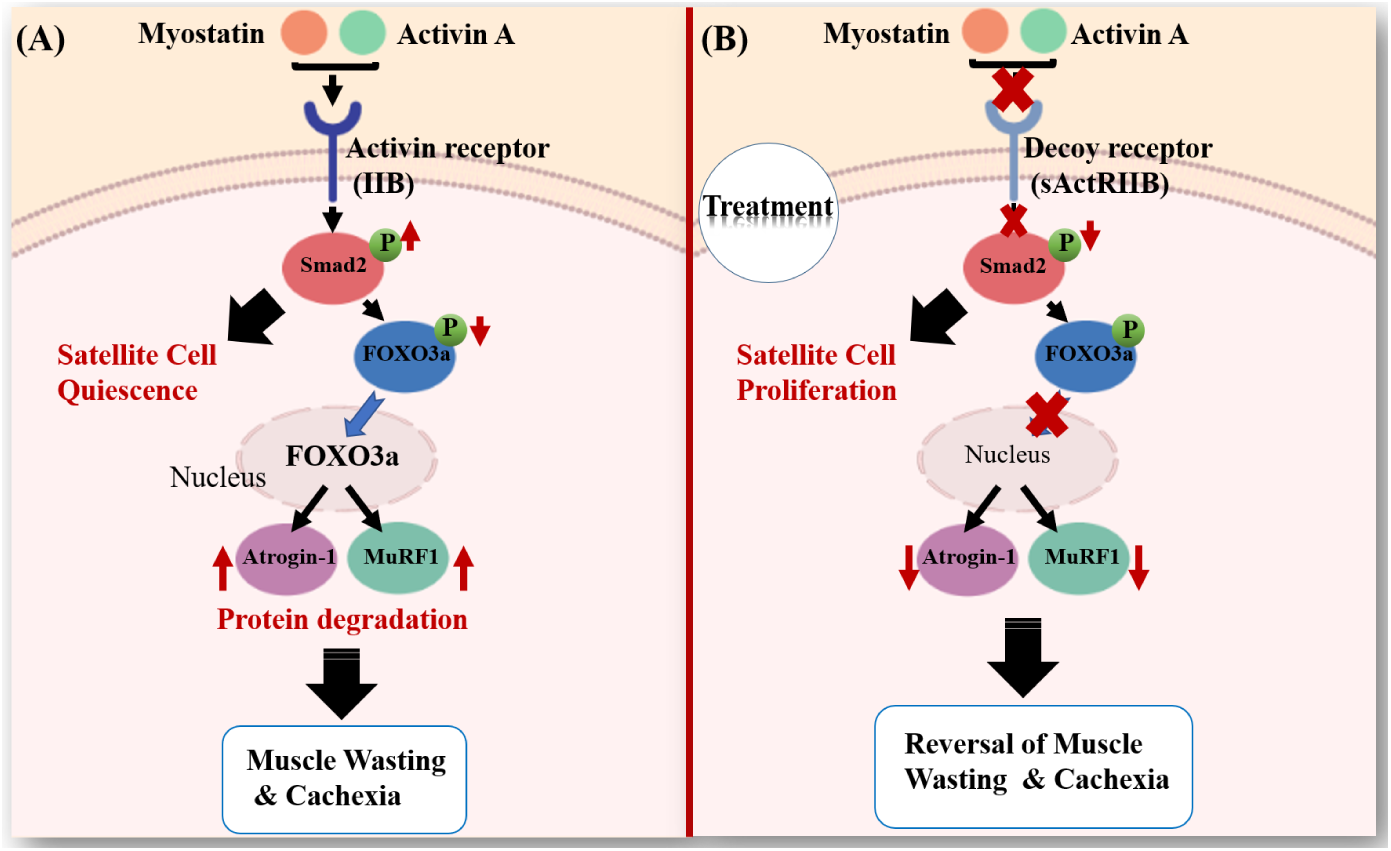
Myostatin and activin signaling pathway in muscle. (A) Myostatin and activin A bind to activin type IIB receptor (ActRIIB) on muscle cell membranes resulting in phosphorylation of Smad2. The phosphorylated Smad2 induces satellite cell quiescence and the dephosphorylation of transcription factor FOXO3a, which consequently moves into the nucleus and activates the transcription of E3 ubiquitin ligase, MuRF1 and Atrogin-1 These muscle-atrophy enzymes provide the specificity that causes the degradation of muscles leading to cachexia. (B) Using a soluble form of the activin type IIB receptor, sActRIIB, the effect of myostatin/activin A signaling is blocked, resulting in a decrease in phosphorylated Smad2, an increase in satellite cell proliferation, and a decrease in ubiquitin enzymes. Blocking this pathway can lead to muscle hypertrophy and reversal of cachexia (35).

#### Healthy Experimental Data

To parameterize the healthy feedback control mechanisms that exist in normal growth and development, we fit our model Eq. (1)–Eq. (3) to average body weight measurements of the US-bred male CDF1 mouse. Growth data is available from the supplier for the first 3 to 15 weeks of life (51) which does not fully characterize the adult mouse size. We thus extend this data to 300 days by fitting a logistic growth curve to the average body weight:

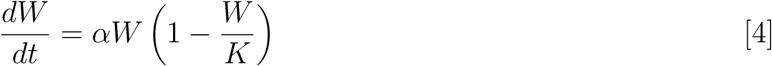

with a growth rate of *α* = 0.0756, a carrying capacity of *K* = 28.3 g, and an initial mass at 3 weeks, day *t* = 0, of *W*(0) = 11.26g. Experimental data points are then extended beyond 15 weeks (84 days), by sampling every 10 days from day 90 up to day 300 from this logistic growth model. Fitting with the extended data ensures the model parameters capture the homeostatic adult muscle mass size. Experimental and extended body weight data converted to lean body mass is shown in Figure 1.

#### Simulated Annealing For Parameter Fitting

The full-feedback model is difficult to parameterize due to the nonlinear structure and high sensitivity around 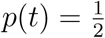. Parameterization is thus performed hierarchically by fixing two model parameters (*d* and *m*) and using a simulated annealing algorithm (52, 53) to fit the remaining parameters (*p*_0_, *p*_1_, *ν*_0_, and *ν*_1_). After a successful fit, parameters *d* and *m* are varied one at a time and simulated annealing is used to find the remaining parameters again. The final fit is determined by the minimum of our objective function, the root-mean-squared-error (RMSE), considering both estimated and extended muscle and satellite mass data. The parameter values determined in this manner are listed in Table (1). The model fit to the data is shown in Figure 1.

## 3. Modeling Cancer Cachexia And Treatment

Myostatin and activin A are two promising molecular targets of muscle wasting in cancer cachexia (18, 54). Myostatin and activin, from the TGF-*β* superfamily, are both upregulated in patients with various chronic diseases such as kidney disease, congestive heart failure, diabetes, sarcopenia, obesity, and advanced stages of cancer (5, 18). They can bind to the activin type IIB receptor (ActRIIB), activating a cascade that leads to satellite cell quiescence and muscle degradation, see Figure 2(A).

Pharmacological blockade of the ActRIIB pathway has been shown to partially-reverse cancer-induced muscle wasting while having no effect on tumour growth or fat loss (21). Using the soluble decoy receptor and inhibiting the Myostatin/activin A signaling increases the amount of phosphorylated FOXO3a and results in attenuated expression of ubiquitin ligases, responsible for muscle degradation and enhanced proliferation of satellite cells, see Figure 2(B). Zhou *et al* (21) showed that this treatment can recover skeletal and heart muscle loss and increase survival times in several mouse models of cancer cachexia. Together with other work on myostatin (55), ActRIIB has been proposed as a promising new treatment approach for cachexia in cancer and other chronic diseases.

### A. Muscle Loss in Cancer Cachexia

Below we summarize the three molecular signaling events that we include in our mathematical formulation believed to contribute to muscle wasting in animal models of cancer cachexia. Cancer cells release systemic signaling factors that travel to muscle tissues through the circulation and activate the ActRIIB pathway. Through this and possibly other pathway activations, the following may occur:

1. satellite cells become quiescent, or have a reduced proliferation rate (10, 35),
2. satellite cells undergo apoptosis, necrosis, or may become permanently quiescent (56–59), and
3. muscle cells atrophy (60).

We modify our model of muscle development, equations Eq. (1)–Eq. (3), to incorporate these new mechanisms resulting from interactions between the tumor and muscle/satellite compartments.

To begin, we model the growing tumor volume, *T*, with an exponential-linear model (61). This choice reflects the fact that our experimental tumor growth data only demonstrates periods of exponential or linear growth(21). Initially, there is a period of exponential growth with a growth rate *µT*. This is followed by a period of linear growth, with growth rate *µ*_1_:

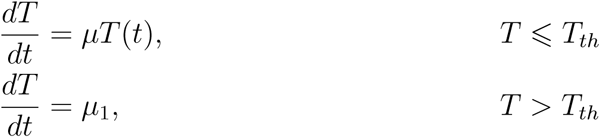

The two growth periods can be connected mathematically, as done in (61), by requiring *µT*_*th*_ = *µ*_1_, so that at the transition point *T*_*th*_, the growth rates are equal. For computational simplicity, the above two growth periods can be combined into one ordinary differential equation, by introducing a transition parameter *η*.

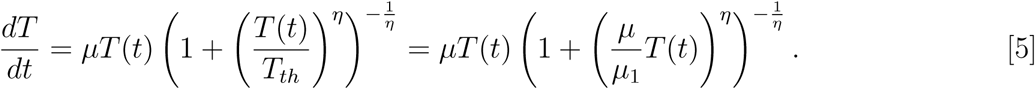

Here *η* controls the speed at which the model transitions from the exponential phase to the linear phase of growth. As done in (61), we use *η* = 20, which is sufficiently fast to capture the transition observed in the data.

Next, we incorporate tumor-induced satellite cell quiescence and interpret this as a suppression of the proliferation rate *ν*(*t*). Thus, cancer reduction of proliferation, which already depends on the size of the muscle mass, is modified by the multiplicative factor 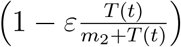. This form allows the strength of the suppression to depend on the tumor volume and can completely suppress proliferation regardless of the current muscle feedback. It also ensures that *ν*(*t*) ≥ 0 for all time. In this suppression factor, *ε* is the maximum reduction possible (with 0 ≤ *ε* ≤ 1) and *m*_2_ is a half-saturation constant for the tumor-size dependent effect.

Finally, we incorporate tumor-induced satellite and muscle cell death by adding terms of the form 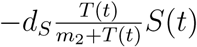 to the satellite cell growth equation, and 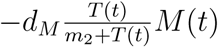 to the muscle cell growth equation. Here *d*_*S*_ and *d*_*M*_ are the tumor-induced satellite cell and muscle cell death rates, respectively, and *m*_2_ is again a half-saturation constant. Modifying our healthy tissue model, equations Eq. (1)–Eq. (3), gives the following mathematical description of the molecular mechanisms involved in cancer cachexia-induced muscle wasting:

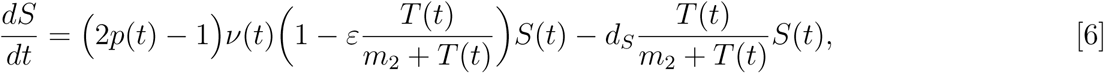

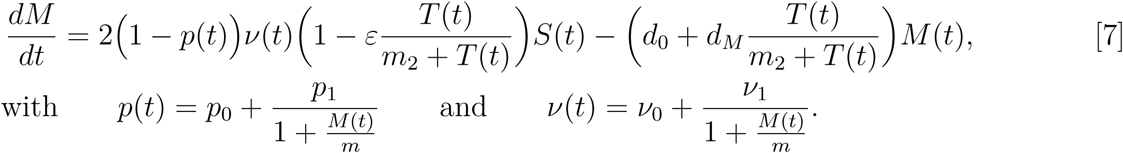

#### Cachexia Model Parameterization

The cachexia mouse model on which we base our parameters is the C26 murine adenocarcinoma model used in (21). Their experimental setup involved subcutaneously injecting 0.5 × 10^6^ C26 cells into the rear flank of 10-week-old male CDF1 mice. We thus assign the initial tumor volume to be *T* (0) = *T*_0_ = 0.5 mm^3^. To determine the growth rate parameters *µ* and *µ*_1_, we minimize the RMSE and fit the numerical solution of Eq. (5) to experimental tumor volume measurements of (21). The result of this fitting is shown in Figure 3(A), with *µ* = 0.446 days^−1^ and *µ*_1_ = 116.0 mm^3^days^−1^. These rates give a transition threshold of *T*_*th*_ = 260 mm^3^, putting the transition from exponential to linear growth around day 14. Exponential-linear tumor growth parameters are listed in Table 2(row 1).

**Table 1.** Parameter values for the healthy muscle model equations Eq. (1)–Eq. (3) found by fitting the model prediction to the estimated and extended muscle and satellite mass data of the CDF1 male mouse. Simulated Annealing was used to find *p*_0_, *p*_1_, *ν*_0_, and *ν*_1_, while *d* and *m* were iterated over manually. The parameters listed correspond to the best fit (minimum RMSE = 0.168). Initial conditions are estimated for a 3-week old CDF1 male mouse.

| Model Parameters |  |  |
| --- | --- | --- |
| $p_0 = 0.479$ | $v_0 = 0.087 \text{ day}^{-1}$ | $d_0 = 0.05 \text{ day}^{-1}$ |
| $p_1 = 0.133$ | $v_1 = 5.591 \text{ day}^{-1}$ | $m = 1000 \text{ mm}^3$ |
| Initial Conditions |  |  |
| $S(0) = 0.536 \text{ g}$ | $M(0) = 10.17784 \text{ g}$ | $\xi = 0.002 \text{ g/mm}^3$ |
| $S(0) = 268.01 \text{ mm}^3$ | $M(0) = 5088.92 \text{ mm}^3$ | |

**Table 2.** Model parameter values and initial conditions. Row 1: Parameter values for the exponential-linear model of tumor growth, Eq. (5), RMSE = 34.11. Row 2: Parameter values for the cachexia model, Eq. (6)–Eq. (7), RMSE=0.157. Row 3-4: initial conditions for experimental groups A and B, respectively, according to Eq. (8). Row 5: Parameter values for cachexia treatment by fitting to experimental group A, Eq. (9)–Eq. (10), RMSE=0.176. Row 6: Parameter values for cachexia treatment by fitting to group B, RMSE=0.317. Parameter values determined by curve fitting the model solution to experimental data from (21) for C26 tumor-bearing mice with or without sActRIIB treatment. Best fit determined by minimizing the root-mean-squared-error between predicted muscle and satellite mass and experimental lean mass data.

| Model Equations | Fitted Parameter Values |  |  |  |
| --- | --- | --- | --- | --- |
| 1. Exp-linear model Eq. (5) | $\mu = 0.446 \text{ days}^{-1}$ | $\mu_1 = 116.0 \text{ mm}^3 \text{ days}^{-1}$ | $\eta = 20$ | $T(0) = 0.5 \text{ mm}^3$ |
| 2. Cancer-Cachexia Eq. (6)–Eq. (7) | $\varepsilon = 0.004$ | $d_S = 0.030 \text{ days}^{-1}$ | $d_M = 0.104 \text{ days}^{-1}$ | $m_2 = 338 \text{ mm}^3$ |
| 3. Group A ICs | $M^A(0) = 4799.25 \text{ mm}^3$ | | $S^A(0) = 252.75 \text{ mm}^3$ | |
| 4. Group B ICs | $M^B(0) = 4768.85 \text{ mm}^3$ | | $S^B(0) = 251.15 \text{ mm}^3$ | |
| 5. sActRIIB Treatment Eq. (9)–Eq. (10) Group A | $A_1 = 0$ | $A_2 = 0$ | $A_3 = 0.51$ | $A_4 = 0.54$ |
| 6. Group B | $A_1 = 0$ | $A_2 = 0$ | $A_3 = 0.62$ | $A_4 = 1$ |

**Fig. 3.**
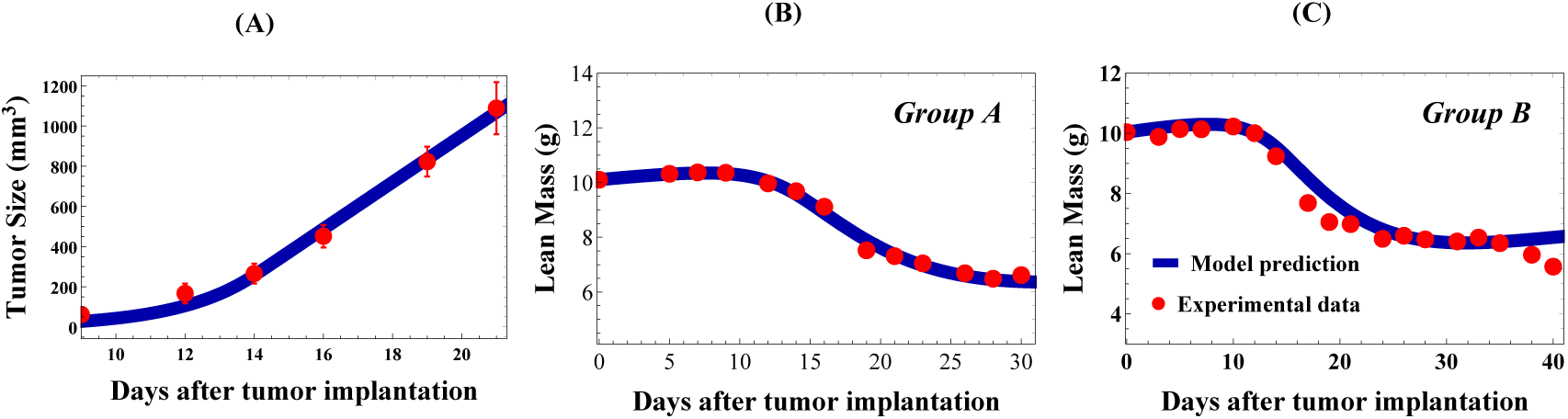
Tumor growth and cachexia model fits to experimental data (21). (A) Exponential-linear tumor growth model (equation Eq. (5), blue curve) fit to C26 experimental tumor volume data from (21) (red points). Initial condition for the numerical simulation is chosen to match experimental conditions, with *T* (0) = *T*_0_ = 0.5 mm^3^ (or 0.5 × 10^6^ cells) representing the initial tumor implant on day 0. Model parameter values are listed in Table 2(row 1). (B) Cachexia model prediction (equations Eq. (5)–Eq. (7), blue curve) fit to lean mass experimental data from mouse group A (red points). In (C), the model prediction is validated by comparing to lean mass experimental data from mouse group B (red points). Cachexia model parameter values are listed in Table 2(row 2), with initial conditions in rows 3 (group A) and row 4 (group B).

To determine the cachexia-related model parameters, *ε, d*_*S*_, *d*_*M*_, and *m*_2_, we fit model predicted lean mass, *ξ*(*M* + *S*), from equations Eq. (5)–Eq. (7) to group A of the C26 mouse experimental data (21). These parameters are determined by curve fitting to minimize the RMSE between the experimental mass data and our model prediction, see Figure 3(B). Resulting cachexia parameter values are listed in Table 2(row 2). As done with the healthy mouse body weight data, we decomposed the cachexia body weight measurements by assuming lean mass corresponds to 40% of the body mass after subtracting predicted tumor mass (based on equation Eq. (5)) and the conversion factor *ξ* = 0.002 g/mm^3^.

Numerically, the system of equations Eq. (5)–Eq. (7) is solved using initial conditions *T* (0) = 0.5 mm^3^, *S*(0) = 252.75 mm^3^, and *M* (0) = 4799.25 mm^3^, and base model parameters from Table 1. The stem and muscle initial conditions correspond to the extrapolated *M* (*t*) and *S*(*t*) values from the body weight on injection day, using the predicted stem ratio from the healthy model, equations Eq. (1)–Eq. (3), solved on day 49 which corresponds to a 10-week old male CDF1 mouse 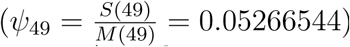. For example, the average body weight of the 10 mice in experimental group A is about 25.26 g. Thus, we start the numerical simulation with

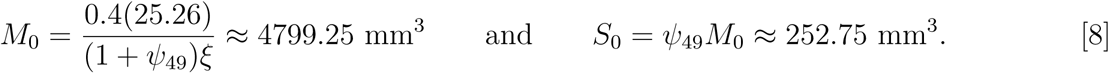

This ensures that the model prediction is fit to experimental data while keeping the developmental stem ratio, and resulting feedback mechanisms, age-appropriate.

Fitting the model to experimental group A from (21) results in parameter values *ε* = 0.004, *m*_2_ = 338 mm^3^, *d*_*S*_ = 0.030 day^−1^, and *d*_*M*_ = 0.104 days^−1^, and fit shown in Figure 3(B). Comparing our model prediction to animal group B, with the same experimental set-up, is shown in Figure 3(C). All parameter values are the same between the two Figures, with the exception of initial conditions for *M* and *S*. These are set according to Eq. (8) with the group’s average lean mass on day 0, 25.10 g for group B. Good agreement is observed between the model prediction and experimental data in both groups.

The model suggests that cachexia requires increased death rates to both muscle and satellite cells. The mechanism of satellite cell quiescence, however, is not required by the model to match experimental observations, as setting *ε* = 0 results in approximately the same fit with the same RMSE, see Figure 7(A). Alternatively, since the mass of the satellite compartment is quite small, the mechanism of quiescence may be appearing in our model as a death rate (permanent quiescence) rather than as a decrease in proliferation frequency.

**Fig. 4.**
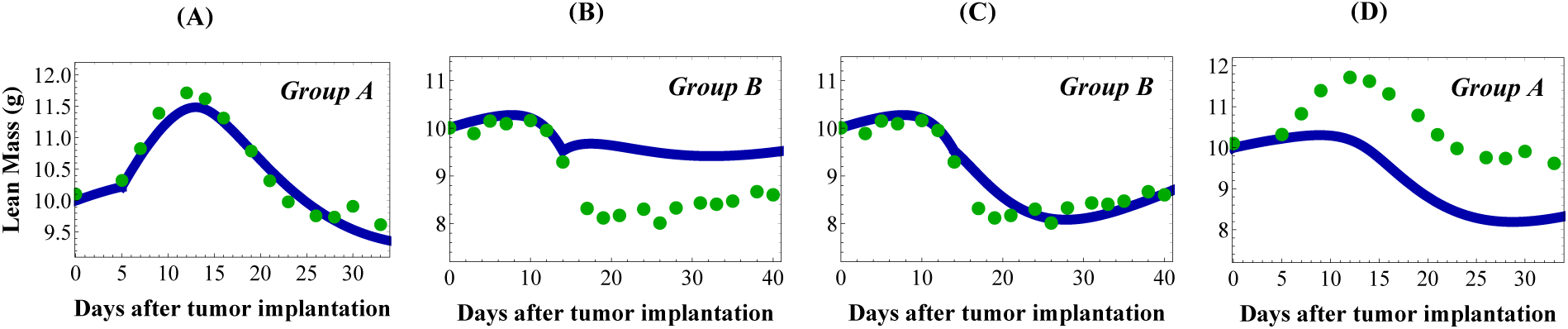
Model prediction of cachexia treatment at either early cachexia (group A, treatment starting on day 5) or late cachexia (group B, treatment starting on day 14). (A) The result of parameter fitting to group A, and (B) the prediction of group B using group A parameter values. (C) The result of parameter fitting to group B, and (D) the prediction of group A using group B parameter values. Experimental data (Green dots) and model simulation (blue line).

**Fig. 5.**
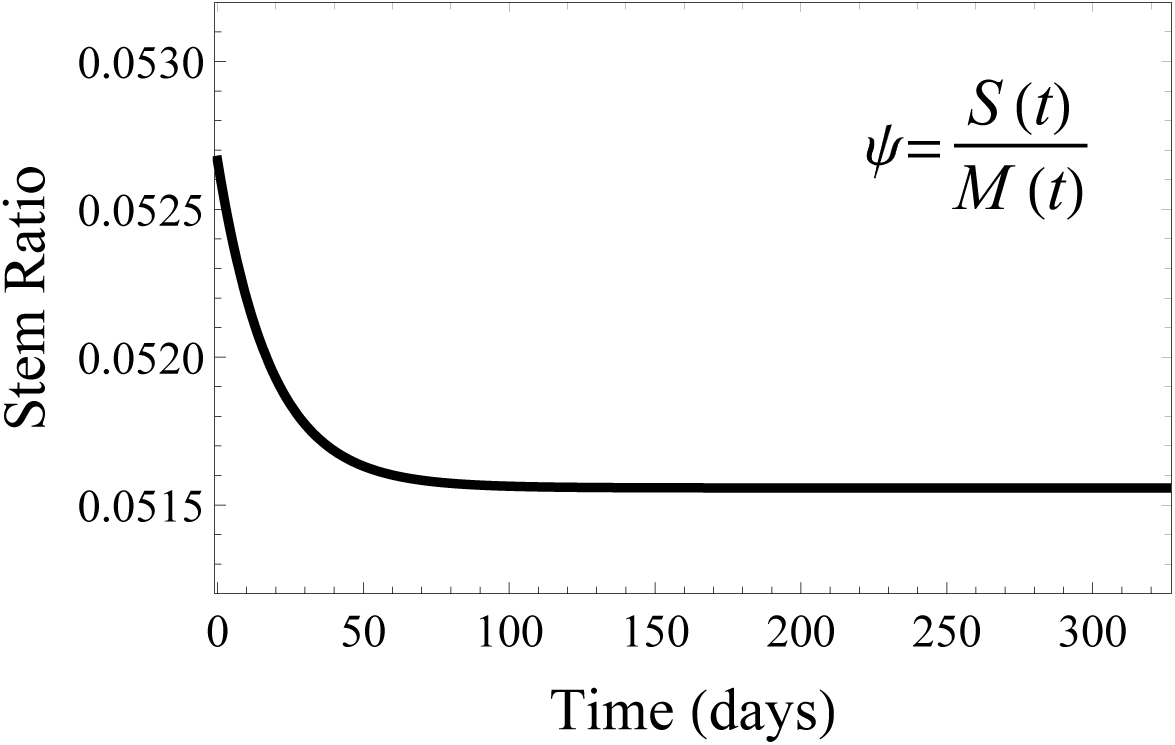
Model simulation of the stem cell ratio *ψ*(*t*) = *S*(*t*)*/M* (*t*). The ratio starts higher due to our estimated initial conditions and the period of growth in the first 3^*rd*^ − 15^*th*^ weeks of life. Once the adult lean mass is obtained, the stem ratio reaches the steady-state homeostatic value.

**Fig. 6.**
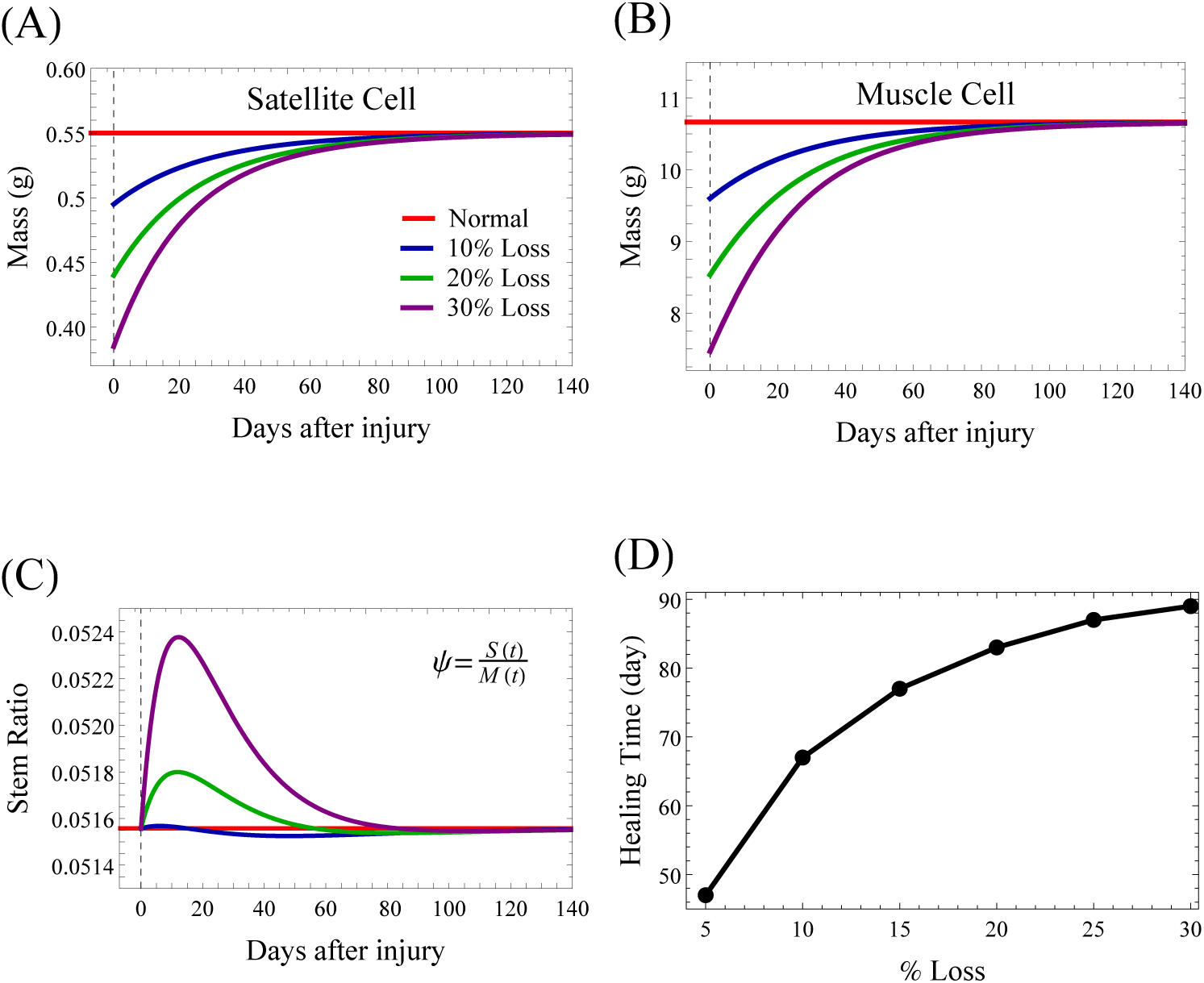
Model prediction of the skeletal muscle response to an injury of 10, 20, or 30% loss in each model compartment. Pre-injury the muscle cells are at the homeostatic level. Post-injury, the satellite cells are dividing to renew themselves (A) and differentiate to new muscle cells (B) and both compartments slowly return to their homeostatic (pre-injury) levels. (C) The stem ratio increases transiently in response to the injury. (D) Healing time for skeletal muscle after a %-loss injury to both muscle and satellite model compartments. Healing time is defined by the time required to re-obtain the pre-injury homeostatic mass computationally as (|*M* (*t*) − *M*_*SS*_ | *<* 0.1).

**Fig. 7.**
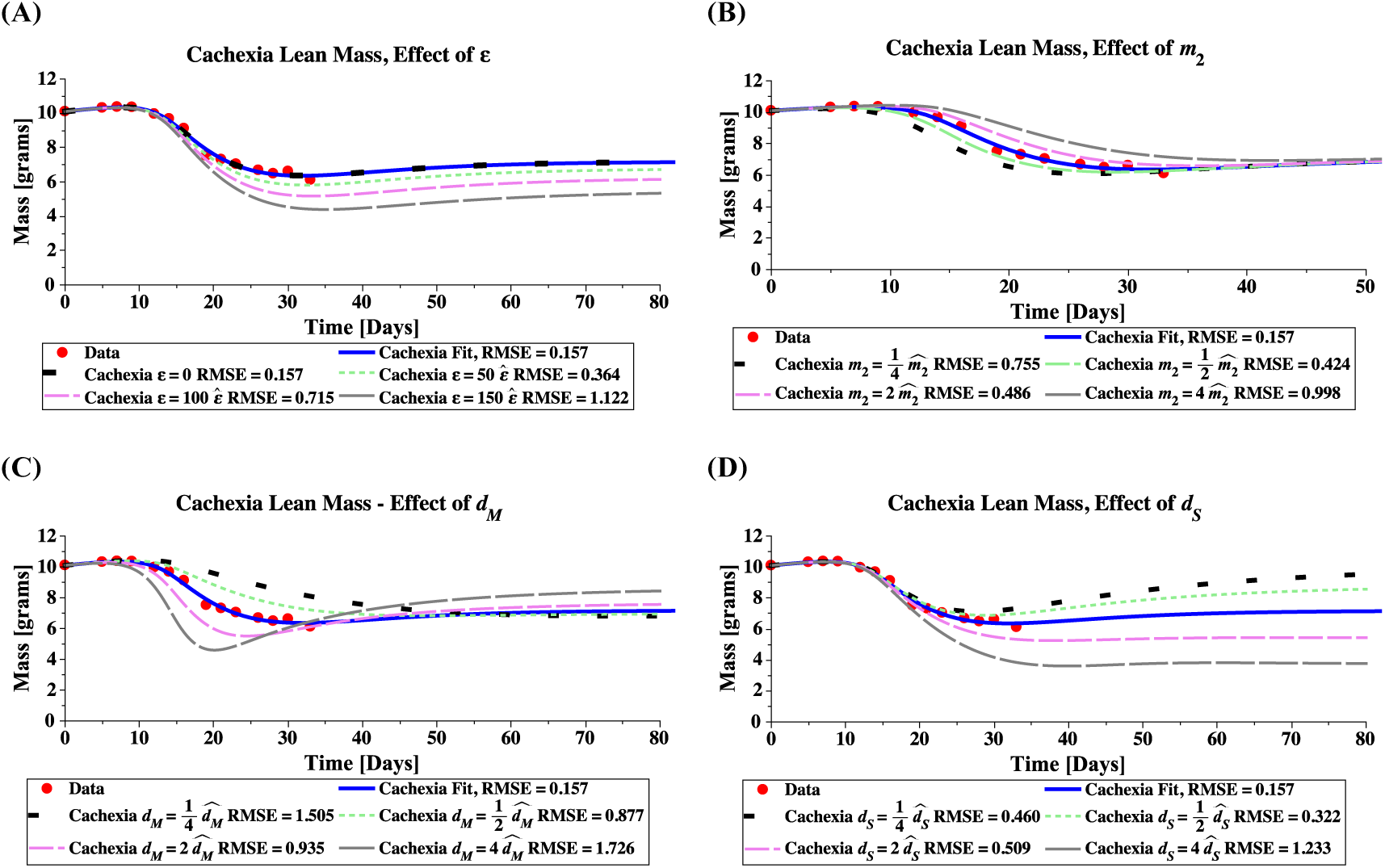
Model simulations of lean mass showing the effect of cachexia parameters *ε* (A), *m*_2_ (B), *d*_*M*_ (C), and *d*_*S*_ (D). Parameter value estimates 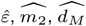, and 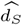, are as listed in Table 2(row 2), unless otherwise specified. All simulations use initial conditions for group A.

### B. Reversing Muscle Loss Through Anti-Cachexia Treatment

Treatment with soluble activin type IIB-receptor (sActRIIB) inhibits myostatin/activin A signaling and leads to dramatic muscle growth in vivo. From Figure 2, we see that two downstream effects of blocking the ActRIIB pathway are satellite cell proliferation and enhanced muscle protein synthesis. Note that, while the treatment seems to reverse muscle wasting, it does not affect tumor growth in the C26 model.

To incorporate sActRIIB treatment into our mathematical model of cancer cachexia, equations Eq. (5)–Eq. (7), we introduce treatment parameters *A*_1_, *A*_2_, and *A*_3_ which reduce the tumor-induced effects of *ε, d*_*S*_, and *d*_*M*_, respectively. Additionally, we introduce treatment parameter *A*_4_ which allows the treatment to reduce the natural muscle death rate *d*_0_. For each treatment parameter we require 0 ≤ *A*_*i*_ ≤ 1, for *i* = 1 … 4. The cachexia-treatment model is thus, equation Eq. (5) (unmodified by sActRIIB treatment) and

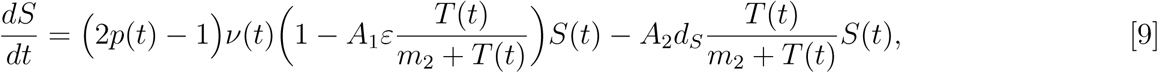

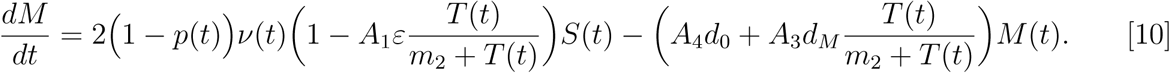

#### Cachexia Treatment Parameterization

Treatment parameters *A*_*i*_ are determined by fitting this model to C26 + sActRIIB mouse data (21). Two experimental groups are presented, one where treatment starts in early cachexia on day 5 (group A), and another where treatment starts in late cachexia on day 14 (group B). To determine model parameters, we again minimize RMSE and find the values reported in Table 2(row 5 and 6), for fits to group A and B respectively.

In both group fits, *A*_1_ = *A*_2_ = 0, suggesting that treatment completely blocks the cancer-imposed stem death rate *d*_*S*_ and the decrease in stem proliferation rate *ε*. Thus, the modeling suggests that treatment reactivates the quiescent satellite cell population. In group A’s fit, we find *A*_3_ = 0.51 and *A*_4_ = 0.54, corresponding to a decrease in both cancer-imposed muscle death rate *d*_*M*_ and the natural muscle death rate *d*_0_. Decreasing the natural death rate allows the model to predict the bump in lean mass observed around day 13 in group A, Figure 4(A). However fitting to group B, which does not present with such an obvious bump Figure 4(C), we find *A*_3_ = 0.62 and *A*_4_ = 1, corresponding to a decrease in the cancer-imposed death rate *d*_*M*_ and no treatment-alteration to the natural death rate *d*_0_.

## 4. Results

### A. The Healthy Muscle Stem Cell Ratio

During the adolescent growth stage, satellite cells proliferate to increase muscle mass. As the muscle matures, satellite cells become inactive and are held in reserve until needed for muscle repair. Skeletal muscle in adulthood is generally stable with the numbers of satellite and muscle cells relatively unchanged. Under these normal adult conditions, our mathematical model gives the following steady-state:

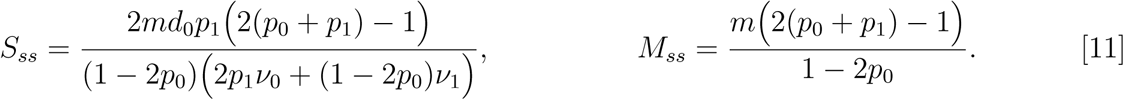

We thus expect the stem cell ratio in the steady state to be

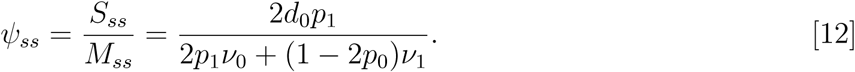

This has a natural upper bound of 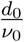, the ratio of the natural muscle cell death rate to the satellite cell homeostatic proliferation rate. From our fitted parameter values in Table 1, the steady-state stem ratio is about *ψ*_*ss*_ = 0.0516. See Figure 5 for the model predicted stem cell ratio for muscle tissue in mice 3 weeks and older.

Sensitivity analysis of the healthy model is performed by computing the relative sensitivity of satellite and muscle compartments to small changes in input parameter values. The homeostatic steady-state, equation 11, is used as the base condition. We then compute the relative change in *S*_*ss*_ and *M*_*ss*_ given a 5% increase or decrease in the parameter values according to the formula:

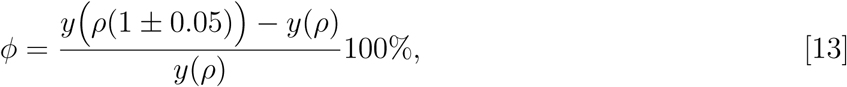

where *y* is either *S*_*ss*_ or *M*_*ss*_, and *ρ* is any model parameter from Table 1. Sensitivity results are summarized in Table 3. The most sensitive model parameter is the homeostatic probability of self-renewing division *p*_0_. As discussed above, when 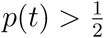 the stem compartment grows without bound, so this result is not unexpected.

**Table 3.**
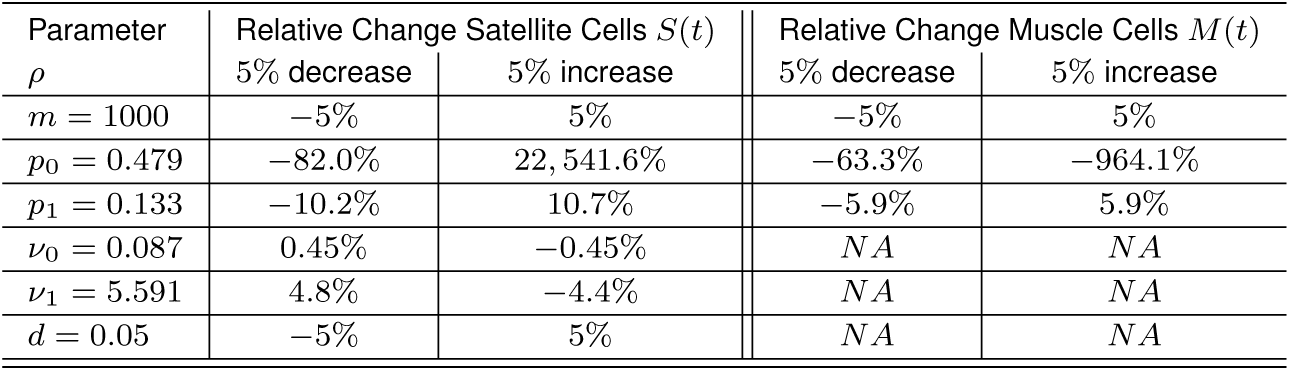
Sensitivity analysis for healthy muscle parameters. The relative change in satellite or muscle compartments is reported given a 5% increase or decrease to the base model parameter value from Table 1. Sensitivity is performed at the steady-state condition, equation 11.

### B. Wound Healing in The Healthy State

After damage, even severe and repeated injuries, skeletal muscle has a remarkable regenerative capacity. Pro-inflammatory signals from the wound area activate satellite cells to proliferate, providing myoblasts to repair muscle fibers.

To test the wound healing ability of our model, we consider three levels of injury: a loss of 10, 20 or 30% in both satellite and muscle cell compartments. Figure 6(A) shows the satellite cell dynamic, which increases to regenerate the muscle, and then slowly returns to its homeostatic (pre-injury) level. As a result of satellite cell proliferation, the number of muscle cells increases until reaching the homeostatic level, Figure 6(B). The number of satellite cells depends on the severity of the lesion, Figure 6(C). The time required to heal the wound depends on the severity of the injury, as shown in Figure 6(D). Larger injuries require longer healing times, according to a sub-linear relationship. The more serious the injury, the longer it takes to re-achieve homeostasis.

### C. Sensitivity Analysis For Cachexia And Treatment

Sensitivity analyses for the cachexia model (Eq. (5)–Eq. (7)) and treatment model (Eq. (5), Eq. (9), Eq. (10)) are now presented. Relative change in muscle or satellite cell populations is reported on the day 20^*th*^ post implant to determine the sensitivity during the dynamic period of muscle loss (rather than the steady-state which is likely not obtained by the cachexic animal). Similar to the above, we iteratively consider a 5% increase or decrease to the cachexia or treatment base parameter values, using Eq. (13) to compute the relative change.

Under cachexia, the top of Table 4, the model is most sensitive to the tumor-induced death rates for satellite cells (*d*_*S*_) and muscle cells (*d*_*M*_). The satellite cell population is most sensitive to *d*_*S*_ as this parameter directly affects *S*(*t*) while the death rate for muscle cells affects the stem population indirectly through the feedback mechanisms. The muscle mass is most sensitive to parameter *d*_*M*_, with *m*_2_ the second most sensitive as this parameter shifts the half-saturation point of the tumor’s influence. Note that since parameter *ε* (tumor-induced quiescence) is very close to zero, it’s sensitivity is also essentially zero. If the base value of *ε* was larger, we would see a more significant sensitivity here (see Figure 7).

**Table 4.** Sensitivity analysis for cachexia and treatment model parameters. The relative change in satellite or muscle compartments is reported given a 5% increase or decrease to the estimated value from Table 2(rows 2 and 5). Sensitivity is performed on day 20 post tumor implantation to capture sensitivity during the dynamical range of interest (during muscle loss).

| Cachexia Parameters |  |  |  |  |
| --- | --- | --- | --- | --- |
| Parameter | Relative Change Satellite Cells $S(20)$ | | Relative Change Muscle Cells $M(20)$ | |
| $\rho$ | 5% decrease | 5% increase | 5% decrease | 5% increase |
| $m_2 = 338$ | -0.074% | 0.074% | -0.71% | 0.69% |
| $\varepsilon = 0.004$ | 0.00 | 0.00 | 0.00 | 0.00 |
| $d_S = 0.030$ | 0.72% | -0.72% | 0.16% | -0.16% |
| $d_M = 0.104$ | -0.46% | 0.46% | 1.55% | -1.51% |
| Treatment Parameters (Fit to Group A) |  |  |  |  |
| parameter | 5% decrease | 5% increase | 5% decrease | 5% increase |
| $A_3 = 0.51$ | -0.15% | 0.15% | 0.95% | -0.94% |
| $A_4 = 0.54$ | -0.35% | 0.36% | 1.11% | -1.09% |

Under treatment, bottom of Table 4, we have left out parameters *A*_1_ and *A*_2_ since they are estimated to have values of zero. We report only sensitivity for parameter *A*_3_, the treatment reduction of tumor-induced muscle death rate, and parameter *A*_4_, the treatment reduction of the natural death rate. We use parameter estimates obtained from fitting to experimental group A, when treatment was given early in cachexia on day 5. Again sensitivity was performed on day 20 post implant to capture the dynamical region of muscle loss and recovery. Both compartments are more sensitive to *A*_4_ (reduction to the natural death rate) than to parameter *A*_3_ (reduction to the tumor-induced death rate) but this difference is likely not significant as their mechanisms of action in the model are similar.

### D. Mechanisms of Cachexia Target Muscle or Satellite Cells With Different Effects

Simulations of the cachexia model, equations Eq. (5)–Eq. (7), demonstrate that the various mechanisms of cachexia (represented by model parameters *ε, m*_2_, *d*_*M*_, and *d*_*S*_) have differing biological implications, as shown in Figure 7. Briefly, targeting muscle cell directly through an increased death rate leads to rapid mass loss that is recovered in time. Whereas targeting satellite cells by inducing quiescence or death, leads to a permanent loss of lean mass.

Increasing parameter *ε* in the satellite cell proliferation suppression factor 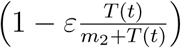 increases the total loss of lean mass, Figure 7(A). According to the assumed negative feedback mechanism that directly affects the proliferation probability and division rate of satellite cells through the function 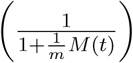 in Eq. (2) & 3, the muscle cell reduction triggers the release of signals that stimulate satellite cell proliferation and differentiation, Figure 8(AI & II). The greater the amount of muscle loss in the body, the stronger the activated feedback signaling, Figure 8(AIII). The feedback proliferation is simultaneously suppressed by the tumor-host interaction factor 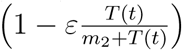, that multiplies the division rate *ν*(*t*) in Eq. (6) & 7 and turns satellite cells quiescent. As *ε* becomes larger, the inhibitory effect on satellite cell proliferation is also larger, see Figure 8(AIV). Therefore the satellite compartment is not able to maintain the healthy steady-state, and as a result mass is lost.

**Fig. 8.**
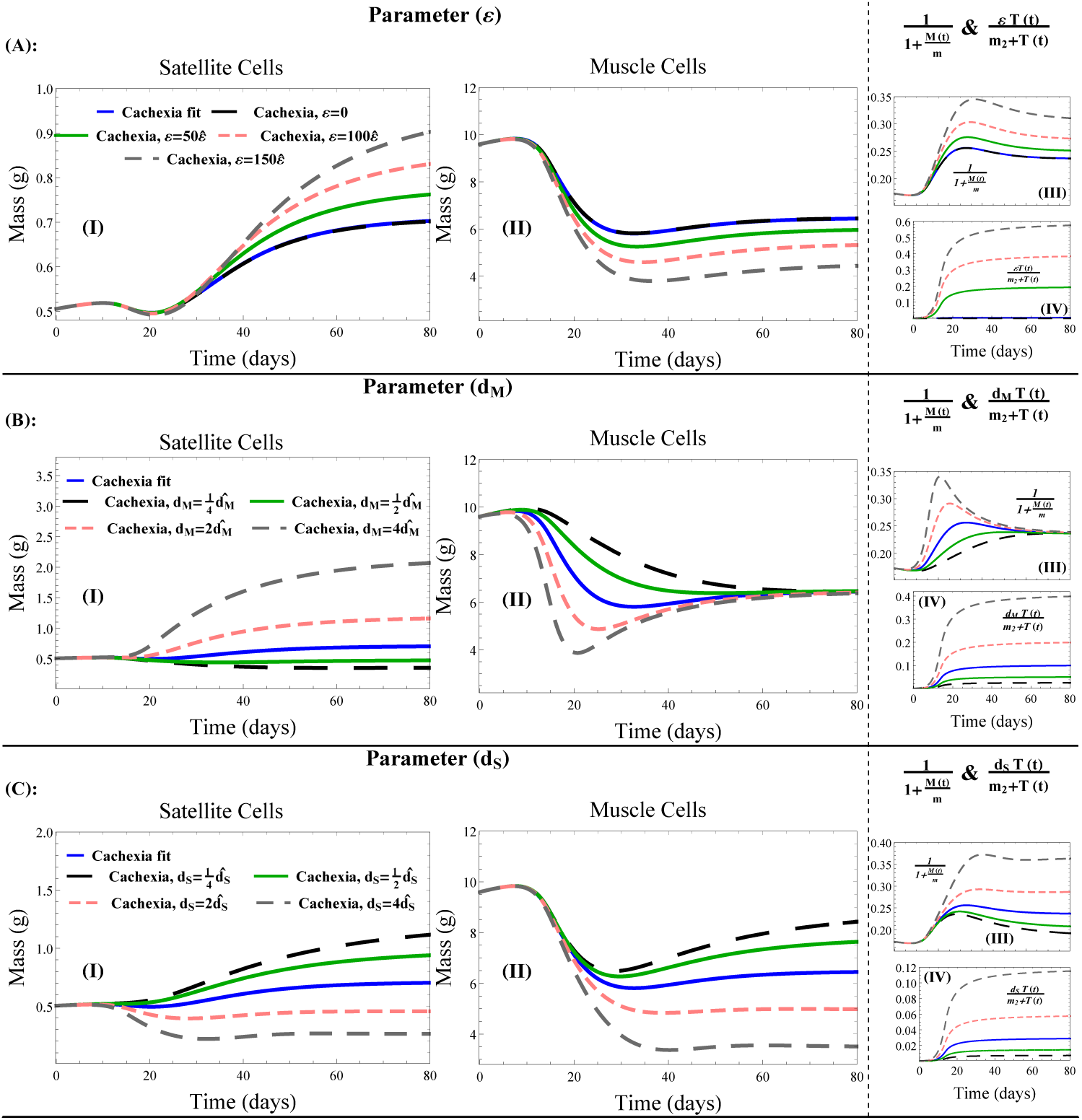
Model simulations of muscle and stem compartments, showing the effect of cachexia parameters *ε* (A), *d*_*M*_ (B), and *d*_*S*_ (C). Parameter value estimates 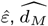 and 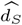, are as listed in Table 2(row 2), unless otherwise specified. All simulations use initial conditions for group A.

Increasing the half saturation constant *m*_2_, delays the transition from health to cachexia, Figure 7(B), as largers tumors are required to induce the same level of signaling.

Increasing parameter *d*_*M*_, the tumor-induced muscle death rate, leads to a rapid loss of muscle mass in the body, Figure 8(BII). As a result, satellite cells are activated by the feedback mechanism, to self renew and create new healthy muscle fibers, Figure 8(BI). The larger the tumor-induced death rate, the larger the feedback effect, Figure 8(BIII). This response by the satellite cells partially recovers the muscle mass loss but requires a significant increase in satellite mass, resulting in a higher steady-state mass, which is likely not realized by the host, Figure 7(C). Importantly, the model predicts that while an increased muscle death rate can lead to rapid mass loss, the effect is not lasting, and mass will be recovered by the stem cells given sufficient time.

Increasing parameter *d*_*S*_, the tumor-induced satellite cell death rate, decreases the stem cell reserve, Figure 8(CI), and results in a reduced muscle cell mass at steady state, Figure 8(CII). As muscle mass drops, the feedback system attempts to activate satellite cells, Figure 8(CIII). The tumor-induced stem death rate is a significant mechanism of lean mass loss that cannot be overcome by feedback activation, Figure 8(CIV). As a result, lean mass steady state decreases with increasing tumor-induced stem death rate, Figure 7(D).

### E. Treatment Partially Restores Muscle Mass by Reactivating Satellite Cells

Using our model of cachexia treatment, we now examine the effects of treatment on lean mass. Figure 9 shows the model prediction using treatment parameter values determined by fitting with either group A, where mice were treated in early cachexia on day 5, or group B, where mice were treated in late cachexia on day 14. Parameter values are as listed in Table 2. In both fits, the treatment blocks cachexia mechanisms targeting satellite cells, resulting in their reactivation, Figure 9(A & D). The mechanisms targeting muscle cells are reduced by about 50% in efficacy. The main difference between the fits is that the natural muscle death rate is reduced in group A (in order to capture the lean mass bump observed post treatment), whereas it is not affected at all in group B, see Figures 9(B & E).

**Fig. 9.**
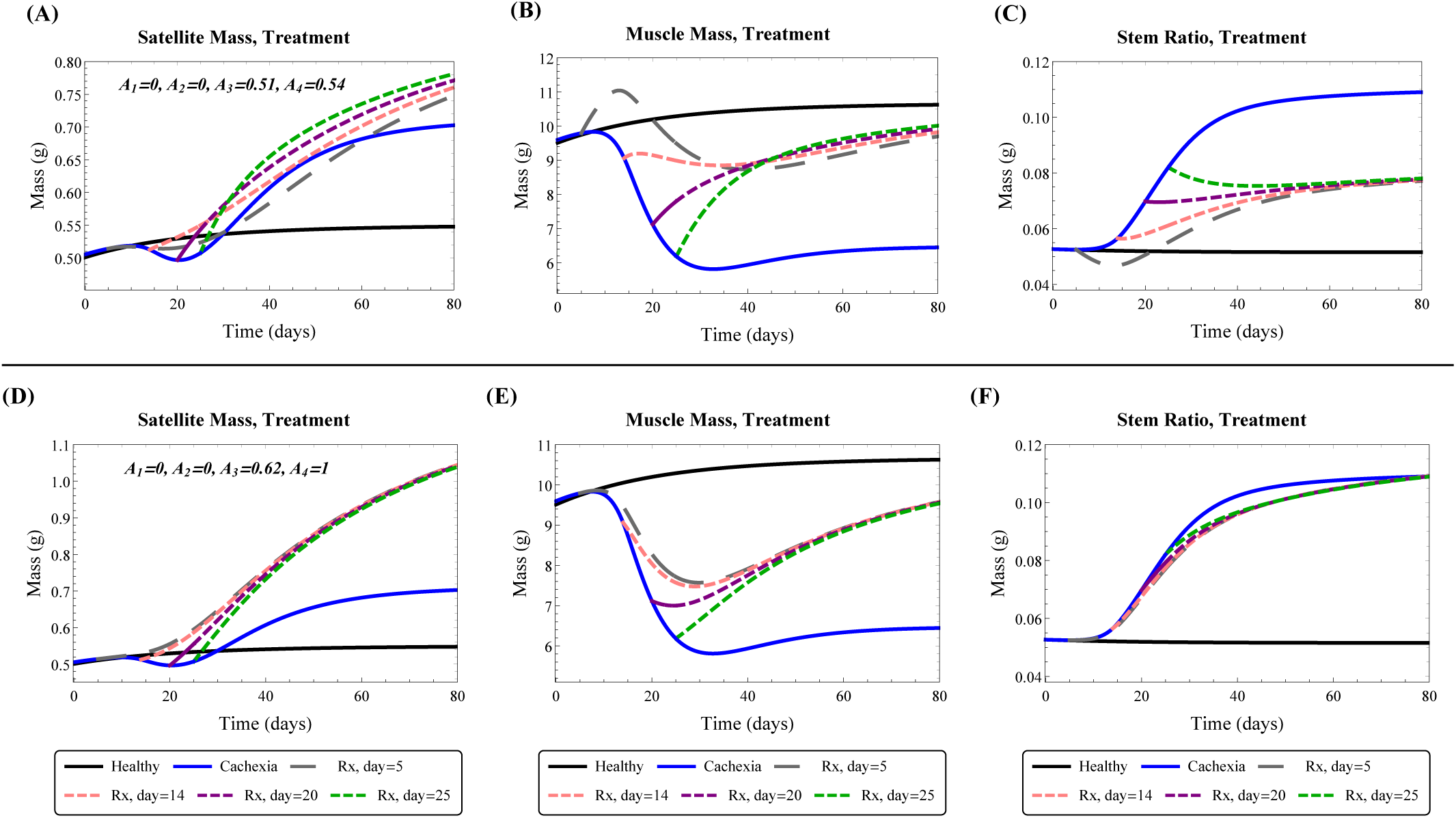
Model predicted satellite cell mass (A,D), muscle mass (B,E), and stem ratio (C,F) under cachexia treatment starting on day 5, 14, 21, or 25 after tumor implantation. Top row (A,B,C) uses treatment parameters from fitting to group A. Bottom row (D,E,F) uses treatment parameters from fitting to group B.

Removal of the inhibition of satellite cells by cachexia treatment results in their reactivation and proliferation to replenish the lost muscle mass. With treatment only partially blocking muscle cell death, the natural feedback pushes the system to obtain a new steady-state where lean mass is below the healthy level and the stem ratio is higher, see Figures 9(C & F). Interestingly, when the natural muscle death rate is reduced by treatment, group A fit, the stem ratio converges to about 8%, while it converges to the predicted cachexia stem ratio of 10% when the natural death rate is unaffected by treatment, group B fit. This is a result of the maximal muscle death rates (natural plus cachexia), which are:

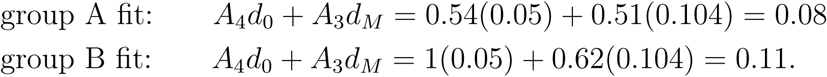

Thus, the model predicts a higher muscle death rate when treatment occurs in late cachexia, than when treatment occurs earlier. Both of these rates are still higher than the natural death rate, which results in a lean mass steady state smaller than in the healthy state.

## 5. Discussion

In this study, we mathematically investigate cancer-associated muscle wasting through the intracellular ActRIIB signaling pathway. Using a stem-hierarchy framework, we present a new model of cancer cachexia and treatment that is parameterized with C26-tumor-bearing CDF1 mouse experimental data. We extend our cancer cachexia model to examine the consequences of anti-ActRIIB treatment causing a partial reversal of muscle loss.

Cachexia is defined as the extreme loss of lean body mass and is associated with multiple diseases including cancer. In studies of cancer cachexia, several signaling pathways have been investigated for their role in regulating muscle atrophy-hypertrophy balance (62, 63). However, it was recently shown that the myostatin/activin-ActRIIB pathway is a dominant pathway leading to muscle degradation through the ubiquitin-proteasome proteolytic pathway. Inhibition of this pathway can reverse muscle loss in different diseases such as Duchenne muscular dystrophy (DMD) (64–67), metabolic diseases (obesity and diabetes) (68, 69), androgen deficiency (70), X-linked myotubular myopathy (XLMTM) (71), amyotrophic lateral sclerosis (ALS) (72), and limb-girdle muscular dystrophy (LGMD) (73).

In this work, we presented a mathematical model describing the complex behavior of satellite cells during skeletal muscle growth and regeneration in response to injury or disease. The model predicts muscle and satellite cell dynamics in the states of health, cancer cachexia, and anti-cachexia treatment. Model predictions demonstrate how the feedback mechanisms of healthy tissue activate to repair injury by stimulating satellite cells to proliferate and differentiate, restoring muscle mass. In cachexia, tumor growth causes muscle mass loss through three main dynamics: (1) reducing satellite cell proliferation rate, (2) inducing death of satellite cells, and (3) inducing death of muscle cells. These negative tumor effects on lean body mass are persistent in that our model of anti-cachexia treatment cannot fully restore the healthy state.

Our model predictions are consistent with previously published and experimentally validated work. In (74), the authors propose that myostatin and activins are capable of binding to both ActRIIA and ActRIIB (activin type II receptors) with different affinities, and that to achieve a strong functional benefit the blockade of both receptors is needed. The experimental data used here assumes only the effect of ActRIIB pathway blockade, and thus so did our modeling work. Future work can extend the model here to consider ActRIIA pathway blockade which may lead to greater muscle regrowth and full recovery of tumor-induced loss. Furthermore, images of muscle fiber cross-sections (21) suggest that muscle atrophy also involves shrinkage of fiber diameters, which is another mechanism of mass loss that was not considered in our cachexia model. This is potentially an important mechanism of mass loss that should be included in the model and is the subject of future work. We speculate that inclusion of this alternate mechanism may help to explain the bump observed in lean mass recovery after early treatment of cachexia which is not present in late treatment.

The modeling work presented here suggests that anti-cachexia treatment can partially recover muscle mass loss by reactivating satellite cells, resulting in a higher than normal stem ratio. The fact that satellite cell reactivation and differentiation was significant to muscle recovery under treatment is consistent with the experimental evidence reported in (21). However, other studies have reported that muscle hypertrophy after pharmacological inhibition of the myostatin pathway occurs without significant activation of satellite cells, and that therefore the satellite cells play little to no role in muscle hypertrophy when induced by this pathway (75–78). Since the satellite cells are primitive cells that regulate muscle homeostasis, they suggest that myostatin and activin A may predominantly signal directly to myofibers. Here, we have focused on the stem cell hierarchy. The distinctive proliferative behaviors of stem cells emerge as a result of the feedback dynamics, and hence muscle regrowth is a direct consequence of stem cell regenerative potential. If these signaling molecules directly alter myofibers, possibly in altering fiber diameter, then as discussed above, this could be explored in an extension of this work. Future work can also explore the effects of other factors or agents that potentially act as intermediates in signal transmission within the stem cell hierarchy.

## ACKNOWLEDGMENTS

This research has been supported by the Ryerson University Faculty of Science Dean’s Research Fund, by the Ryerson University Department of Mathematics, and by the NSERC Discovery Grant program.

## References

1. Tisdale MJ (2002) Cachexia in cancer patients. Nature Reviews Cancer 2(11):862.

2. Springer J, Von Haehling S, Anker SD (2006) The need for a standardized definition for cachexia in chronic illness. Nature clinical practice Endocrinology & metabolism 2(8):416–417.

3. Evans WJ, et al. (2008) Cachexia: a new definition. Clinical nutrition 27(6):793–799.

4. Donohoe CL, Ryan AM, Reynolds JV (2011) Cancer cachexia: mechanisms and clinical implications. Gastroenterology research and practice 2011.

5. von Haehling S, Anker SD (2010) Cachexia as a major underestimated and unmet medical need: facts and numbers.

6. Dewys WD, et al. (1980) Prognostic effect of weight loss prior tochemotherapy in cancer patients. The American journal of medicine 69(4):491–497.

7. Fearon KC, Voss AC, Hustead DS (2006) Definition of cancer cachexia: effect of weight loss, reduced food intake, and systemic inflammation on functional status and prognosis. The American journal of clinical nutrition 83(6):1345–1350.

8. Tisdale MJ (2009) Mechanisms of cancer cachexia. Physiological reviews 89(2):381–410.

9. Kern K, Norton J (1988) Cancer cachexia. Journal of Parenteral and Enteral Nutrition 12(3):286–298.

10. Mathew SJ (2011) Inactivating cancer cachexia. Disease models & mechanisms 4(3):283–285.

11. Lawson DH, Richmond A, Nixon DW, Rudman D (1982) Metabolic approaches to cancer cachexia. Annual review of nutrition 2(1):277–301.

12. Fearon K, et al. (2011) Definition and classification of cancer cachexia: an international consensus. The lancet oncology 12(5):489–495.

13. Aoyagi T, Terracina KP, Raza A, Matsubara H, Takabe K (2015) Cancer cachexia, mechanism and treatment. World journal of gastrointestinal oncology 7(4):17.

14. Bruggeman AR, et al. (2016) Cancer cachexia: beyond weight loss. Journal of oncology practice 12(11):1163–1171.

15. Del Fabbro E, Inui A, Strasser F (2012) Overview of cancer cachexia in Cancer Cachexia. (Springer), pp. 1–5.

16. Tazi E, Errihani H (2010) Treatment of cachexia in oncology. Indian journal of palliative care 16(3):129.

17. Naito T (2019) Emerging treatment options for cancer-associated cachexia: A literature review. Therapeutics and clinical risk management 15:1253.

18. Han H, Zhou X, Mitch WE, Goldberg AL (2013) Myostatin/activin pathway antagonism: molecular basis and therapeutic potential. The international journal of biochemistry & cell biology 45(10):2333–2347.

19. Sandri M, et al. (2004) Foxo transcription factors induce the atrophy-related ubiquitin ligase atrogin-1 and cause skeletal muscle atrophy. Cell 117(3):399–412.

20. Glass DJ (2005) Skeletal muscle hypertrophy and atrophy signaling pathways. The international journal of biochemistry & cell biology 37(10):1974–1984.

21. Zhou X, et al. (2010) Reversal of cancer cachexia and muscle wasting by actriib antagonism leads to prolonged survival. Cell 142(4):531–543.

22. Elkina Y, von Haehling S, Anker SD, Springer J (2011) The role of myostatin in muscle wasting: an overview. Journal of cachexia, sarcopenia and muscle 2(3):143.

23. Han H, Mitch WE (2011) Targeting the myostatin signaling pathway to treat muscle wasting diseases. Current opinion in supportive and palliative care 5(4):334.

24. McCroskery S, Thomas M, Maxwell L, Sharma M, Kambadur R (2003) Myostatin negatively regulates satellite cell activation and self-renewal. The Journal of cell biology 162(6):1135–1147.

25. Klimek MEB, et al. (2010) Acute inhibition of myostatin-family proteins preserves skeletal muscle in mouse models of cancer cachexia. Biochemical and biophysical research communications 391(3):1548–1554.

26. Murphy KT, et al. (2010) Antibody-directed myostatin inhibition in 21-mo-old mice reveals novel roles for myostatin signaling in skeletal muscle structure and function. The FASEB Journal 24(11):4433–4442.

27. Busquets S, et al. (2012) Myostatin blockage using actriib antagonism in mice bearing the lewis lung carcinoma results in the improvement of muscle wasting and physical performance. Journal of cachexia, sarcopenia and muscle 3(1):37–43.

28. Mathews LS, Vale WW (1991) Expression cloning of an activin receptor, a predicted transmembrane serine kinase. Cell 65(6):973–982.

29. Attisano L, Wrana JL, Cheifetz S, Massague J (1992) Novel activin receptors: distinct genes and alternative mrna splicing generate a repertoire of serine/threonine kinase receptors. Cell 68(1):97–108.

30. Oh SP, Li E (1997) The signaling pathway mediated by the type iib activin receptor controls axial patterning and lateral asymmetry in the mouse. Genes & development 11(14):1812–1826.

31. Lee SJ, McPherron AC (2001) Regulation of myostatin activity and muscle growth. Proceedings of the National Academy of Sciences 98(16):9306–9311.

32. Mammucari C, et al. (2007) Foxo3 controls autophagy in skeletal muscle in vivo. Cell metabolism 6(6):458–471.

33. Zhao J, et al. (2007) Foxo3 coordinately activates protein degradation by the autophagic/lysosomal and proteasomal pathways in atrophying muscle cells. Cell metabolism 6(6):472–483.

34. Cohen S, et al. (2009) During muscle atrophy, thick, but not thin, filament components are degraded by murf1-dependent ubiquitylation. The Journal of cell biology 185(6):1083–1095.

35. Tisdale MJ (2010) Reversing cachexia. Cell 142(4):511–512.

36. Rodriguez-Brenes IA, Komarova NL, Wodarz D (2011) Evolutionary dynamics of feedback escape and the development of stem-cell–driven cancers. Proceedings of the National Academy of Sciences 108(47):18983–18988.

37. Janssen I, Heymsfield SB, Wang Z, Ross R (2000) Skeletal muscle mass and distribution in 468 men and women aged 18–88 yr. Journal of applied physiology 89(1):81–88.

38. Mauro A (1961) Satellite cell of skeletal muscle fibers. The Journal of biophysical and biochemical cytology 9(2):493.

39. Allbrook D, Han M, Hellmuth A (1971) Population of muscle satellite cells in relation to age and mitotic activity. Pathology 3(3):233–243.

40. Hellmuth A, Allbrook D (1971) Muscle satellite cell numbers during the postnatal period. Journal of anatomy 110(Pt 3):503–503.

41. Schultz E (1974) A quantitative study of the satellite cell population in postnatal mouse lumbrical muscle. The Anatomical Record 180(4):589–595.

42. Rudnicki M, Le Grand F, McKinnell I, Kuang S (2008) The molecular regulation of muscle stem cell function in Cold Spring Harbor symposia on quantitative biology. (Cold Spring Harbor Laboratory Press), Vol. 73, pp. 323–331.

43. Snow MH (1977) Myogenic cell formation in regenerating rat skeletal muscle injured by mincing ii. an autoradiographic study. The Anatomical Record 188(2):201–217.

44. Bischoff R (1986) A satellite cell mitogen from crushed adult muscle. Developmental biology 115(1):140–147.

45. Schultz E (1978) Changes in the satellite cells of growing muscle following denervation. The Anatomical Record 190(2):299–311.

46. Dhawan J, Rando TA (2005) Stem cells in postnatal myogenesis: molecular mechanisms of satellite cell quiescence, activation and replenishment. Trends in cell biology 15(12):666–673.

47. Pallafacchina G, Blaauw B, Schiaffino S (2013) Role of satellite cells in muscle growth and maintenance of muscle mass. Nutrition, Metabolism and Cardiovascular Diseases 23:S12–S18.

48. Yin H, Price F, Rudnicki MA (2013) Satellite cells and the muscle stem cell niche. Physiological reviews 93(1):23–67.

49. Lander AD, Gokoffski KK, Wan FY, Nie Q, Calof AL (2009) Cell lineages and the logic of proliferative control. PLoS biology 7(1):e1000015.

50. Kuang S, Gillespie MA, Rudnicki MA (2008) Niche regulation of muscle satellite cell self-renewal and differentiation. Cell stem cell 2(1):22–31.

51. (2019) (https://www.criver.com/products-services/find-model/cd2f1-cdf1-mouse).

52. Corana A, Marchesi M, Martini C, Ridella S (1987) Minimizing multimodal functions of continuous variables with the “simulated annealing” algorithm corrigenda for this article is available here. ACM Transactions on Mathematical Software (TOMS) 13(3):262–280.

53. Bertsimas D, Tsitsiklis J, et al. (1993) Simulated annealing. Statistical science 8(1):10–15.

54. Yoshida T, Delafontaine P (2015) Mechanisms of cachexia in chronic disease states. The American journal of the medical sciences 350(4):250–256.

55. Bondulich MK, et al. (2017) Myostatin inhibition prevents skeletal muscle pathophysiology in huntington’s disease mice. Scientific reports 7(1):14275.

56. Jejurikar SS, Marcelo CL, Kuzon JW (2002) Skeletal muscle denervation increases satellite cell susceptibility to apoptosis. Plastic and reconstructive surgery 110(1):160–168.

57. Jejurikar SS, Kuzon WM (2003) Satellite cell depletion in degenerative skeletal muscle. Apoptosis 8(6):573–578.

58. Ferreira R, et al. (2006) Skeletal muscle atrophy increases cell proliferation in mice gastrocnemius during the first week of hindlimb suspension. European journal of applied physiology 97(3):340–346.

59. Guo BS, Cheung KK, Yeung SS, Zhang BT, Yeung EW (2012) Electrical stimulation influences satellite cell proliferation and apoptosis in unloading-induced muscle atrophy in mice. PloS one 7(1).

60. Talbert EE, Guttridge DC (2016) Impaired regeneration: a role for the muscle microenvironment in cancer cachexia in Seminars in cell & developmental biology. (Elsevier), Vol. 54, pp. 82–91.

61. Simeoni M, et al. (2004) Predictive pharmacokinetic-pharmacodynamic modeling of tumor growth kinetics in xenograft models after administration of anticancer agents. Cancer research 64(3):1094–1101.

62. Stitt TN, et al. (2004) The igf-1/pi3k/akt pathway prevents expression of muscle atrophy-induced ubiquitin ligases by inhibiting foxo transcription factors. Molecular cell 14(3):395–403.

63. Schmierer B, Hill CS (2007) Tgf*β*–smad signal transduction: molecular specificity and functional flexibility. Nature reviews Molecular cell biology 8(12):970–982.

64. Li ZB, Zhang J, Wagner KR (2012) Inhibition of myostatin reverses muscle fibrosis through apoptosis. Journal of cell science 125(17):3957–3965.

65. George Carlson C, et al. (2011) Soluble activin receptor type iib increases forward pulling tension in the mdx mouse. Muscle & nerve 43(5):694–699.

66. Morine KJ, et al. (2010) Activin iib receptor blockade attenuates dystrophic pathology in a mouse model of duchenne muscular dystrophy. Muscle & nerve 42(5):722–730.

67. Pistilli EE, et al. (2011) Targeting the activin type iib receptor to improve muscle mass and function in the mdx mouse model of duchenne muscular dystrophy. The American journal of pathology 178(3):1287–1297.

68. Akpan I, et al. (2009) The effects of a soluble activin type iib receptor on obesity and insulin sensitivity. International journal of obesity 33(11):1265–1273.

69. Zhang C, et al. (2012) Inhibition of myostatin protects against diet-induced obesity by enhancing fatty acid oxidation and promoting a brown adipose phenotype in mice. Diabetologia 55(1):183–193.

70. Koncarevic A, et al. (2010) A soluble activin receptor type iib prevents the effects of androgen deprivation on body composition and bone health. Endocrinology 151(9):4289–4300.

71. Lawlor MW, et al. (2011) Inhibition of activin receptor type iib increases strength and lifespan in myotubularin-deficient mice. The American journal of pathology 178(2):784–793.

72. Morrison BM, et al. (2009) A soluble activin type iib receptor improves function in a mouse model of amyotrophic lateral sclerosis. Experimental neurology 217(2):258–268.

73. Ohsawa Y, et al. (2006) Muscular atrophy of caveolin-3–deficient mice is rescued by myostatin inhibition. The Journal of clinical investigation 116(11):2924–2934.

74. Morvan F, et al. (2017) Blockade of activin type ii receptors with a dual anti-actriia/iib antibody is critical to promote maximal skeletal muscle hypertrophy. Proceedings of the National Academy of Sciences 114(47):12448–12453.

75. Sartori R, et al. (2009) Smad2 and 3 transcription factors control muscle mass in adulthood. American journal of physiology-cell physiology 296(6):C1248–C1257.

76. Amthor H, et al. (2009) Muscle hypertrophy driven by myostatin blockade does not require stem/precursor-cell activity. Proceedings of the National Academy of Sciences 106(18):7479–7484.

77. Wang Q, McPherron AC (2012) Myostatin inhibition induces muscle fibre hypertrophy prior to satellite cell activation. The Journal of physiology 590(9):2151–2165.

78. Lee SJ, et al. (2012) Role of satellite cells versus myofibers in muscle hypertrophy induced by inhibition of the myostatin/activin signaling pathway. Proceedings of the National Academy of Sciences 109(35):E2353–E2360.

